# Genome-wide analyses of transcription factors and co-regulators across seven cohorts identified reduced PPARGC1A expression as a driver of prostate cancer progression

**DOI:** 10.1101/2020.05.12.091066

**Authors:** Manjunath Siddappa, Sajad A. Wani, Mark D. Long, Damien A Leach, Ewy A. Mathé, Charlotte L. Bevan, Moray J. Campbell

## Abstract

Defining altered transcription factor (TF) and coregulators that are oncogenic drivers remains a challenge, in part because of the multitude of TFs and coregulators. We addressed this challenge by using bootstrap approaches to test how expression, copy number alterations or mutation of TFs (n = 2662), coactivators (COA; n= 766); corepressor (COR; n = 599); mixed function coregulators (MIXED; n = 511) varied across seven prostate cancer (PCa) cohorts (three of localized and four advanced disease). COAS, CORS, MIXED and TFs all displayed significant down-regulated expression (q.value < 0.1) and correlated with protein expression (ρ 0.4 to 0.55). Stringent expression filtering identified commonly altered TFs and coregulators including well-established (e.g. ERG) and underexplored (e.g. *PPARGC1A*, encodes PGC1α) in localized PCa. Reduced *PPARGC1A* expression significantly associated with worse disease-free survival in two cohorts of localized PCa. Stable PGC1α knockdown in LNCaP cells increased growth rates and invasiveness and RNA-Seq revealed a profound basal impact on gene expression (~2300 genes; FDR < 0.05, logFC > 1.5), but only modestly impacted PPARγ responses. GSEA analyses of the PGC1α transcriptome revealed that it significantly altered the AR-dependent transcriptome, and was enriched for epigenetic modifiers. PGC1α-dependent genes were overlapped with PGC1α-ChIP-Seq genes and significantly associated in TCGA with higher grade tumors and worse disease-free survival. Together these data demonstrate an approach to identify cancer-driver coregulators in cancer and that PGC1α expression is clinically significant yet underexplored coregulator in aggressive early stage PCa.

## INTRODUCTION

Prostate cancer (PCa) is a high-profile hormone-responsive cancer that displays uncertain progression risks. For men with localized PCa, it is unclear which patients will experience treatment failure following either surgery or radiotherapy. In advanced PCa, it’s similarly unclear which patients will experience a sustained response to androgen deprivation therapy (ADT), and who will experience treatment failure giving rise to ADT-recurrent PCa (ADT-RPCa).

Many of these prognostic uncertainties arise due to changes in the control of androgen receptor (AR) signaling. In the normal prostate gland the AR functions in a large and dynamic multimeric complex to regulate gene expression programs that control tissue homeostasis and epithelial cell fates ^27, 34^. In PCa there are strong examples of how signaling is distorted due to changes in AR expression and structural variation, alongside changes in the members of the AR complex ^9, 15, 26, 28, 35, 47, 58, 59, 64, 86, 93^.

Furthermore, epigenomic events^2, 25, 33, 57, 96^ alter enhancer accessibility of downstream AR target genes and even upstream of the AR gene itself^79^. As a result of these changes to the AR complex and allied epigenetic events, the normal differentiation driving and growth limiting functions of the AR are attenuated and there is enhanced regulation of genes that promote aggressive cancer phenotypes^3, 87, 92, 63^.

This re-wiring of AR signaling is mirrored by disrupted functions of other members of the nuclear hormone receptor super-family^35, 50^, and probably occurs more widely to alter how transcription factors (TFs) function. Re-wiring of TFs changes the consequences of cell signaling and contributes to changes in lineage plasticity that are also associated with aggressive PCa^8, 40, 72^. Therefore understanding how TF signaling occurs in PCa progression can lead to the development of strategies for targeting these changes through targeted epigenetic therapies^6, 21, 42^, or the downstream gene networks may be uniquely drug-sensitive^56, 68^. Furthermore disease-specific enhancers that are newly activated provide rationale for targeted deep sequencing to dissect the interactions of germline and structural variation at tumor-driving enhancers^60, 75^. These studies have revealed a number of high profile and common alterations in TFs and coregulators in PCa. For example, ERG^61^ and NR3C1/GR^39^ have been identified as commonly altered, whereas others such as the PPARγ coactivator PGC1α^88, 89^ are relatively under-investigated. However, given the multitude of TFs and coregulators, many have not been investigated at all in the context of PCa.

The Cancer Genome Atlas (TCGA) and other consortia^13, 23, 74^ established mutation, copy number variation and gene/protein expression in hundreds of tumors from both early and advanced PCa along-side clinical features and patient outcome data. These studies have revealed powerful insights into PCa drivers, and offer up the possibility for secondary analyses focusing specifically on TFs and coregulators. In the current study TCGA and other genome-wide data were used specifically to test how TFs and their co-regulators are altered and associate with PCa outcome.

We utilized seven PCa cohorts; three cohorts of localized PCa (TCGA^10^, MSKCC^81^, OICR^74^) and four cohorts of advanced PCa (MICH^41^ FHCRC^41^, BELT^7^ SU2C^1,66^). In these cohorts we examined the expression and genetic changes in different classes of coregulators and TFs, and within each class identified the most significant and clinically-relevant gene changes. These approaches established the class-specific significant mRNA and protein down-regulation of TFs and co-regulators in localized PCa, which were not further altered in advanced PCa, and are not altered significantly by either mutation or copy number alterations (CNA).

These approaches revealed that *PPARGC1A* (encodes PGC1α) was commonly down-regulated associated with worse diseae outcome in local PCa. Stable knockdown of PGC1α in LNCaP cells increased proliferation, led to a more invasive phenotype, and profoundly changed gene expression patterns both positively and negatively. Combining these data with differentially regulated genes between tumors with high or low *PPARGC1A* expression alongside PGC1α ChIP-Seq data identified a network of 60 genes that are both significantly bound and regulated by PGC1α, and significantly associated with higher Gleason Grade tumors. In parallel, these approaches establish a relatively generic work-flow for analyses of gene families or functional groupings to allow investigators to identify clinically-significant relationships which in turn support functional analyses.

## MATERIALS AND METHODS

### Data analyses, integration and code availability

PCa data was downloaded from cBioPortal^13^.

Analyses was undertaken using R (version 3.6.2) ^43 43 43.^

### Cell culture and materials

LNCaP cells (ATCC) were confirmed to be mycoplasma free everyon month and authenticated using STR profiling. Cells were treated with 5,8,11,14-Eicosatetraynoic acid (ETYA, Calbiochem).

### Cell viability and Scratch Assays

Cellular viability was measured by bioluminescent detection of ATP (CellTitre-Glo^®^ assay kit (Promega)). Cells were plated at 5×10^3^ cells per well in 96-well, white-walled plates, allowed to adhere and treated with ETYA (10 μM) or EtOH (vehicle control) to a final volume of 100 μl for 96 hr. Each experiment was performed in triplicate in triplicate wells. For scratch assay, cells were seeded at 1×10^6^ cells per well in a 6-well plate, allowed to adhere for 48 hr to a confluence of about 80% and then wounded by scratching with p200 sterile pipette tip. The debris were removed, and cells washed to make ensure the edges were smoothed with the same dimensions for experimental and control cells. Cells were incubated, and cell migration was assessed by monolayer gap closure after 48 hr.

### Stable knockdown of PGC1α in LNCaP Cells

PGC1α (encoded by PPARGC1A) was knocked-down with shRNA constructs (TG310260, Origene) in LNCaP cells (LNCaP_shPGC1A) or scrambled shRNA (LNCaP_shCtrl). Two PGC1A targeting constructs (shPGC1A-34 and shPGC1A-35) were selected and maintained in media containing puromycin (0.2ug/ml).

### Western Immunoblotting

Total cellular protein was isolated from exponentially growing cells and lysed in ice cold RIPA buffer containing 1x cOmplete Mini Protease Inhibitor Tablets (Roche). Protein concentration were quantified and 75 ug resolved (SDS-PAGE) using 10% polyacrylamide gels, transferred to PVDF membrane and probed with primary antibody against PGC1α (PA5-72948, Invitrogen) and Beta-Actin (ab8229, abcam) overnight at 4°C. Primary antibody was detected with HRP-linked goat anti-rabbit IgG (abcam) and signal captured (ChemiDoc XRS+ system (Bio-Rad)).

### Unique lists of annotated transcription factors and co-regulators

A comprehensive list of TFs and co-regulators genes was developed by text-mining^54^ Gene Ontology (GO) terms that contained phrases including “positive control of transcription”, “negative control of transcription”, “co-activator”, “co-repressor”. From these GO terms, the HGNC gene name and ENSEMBL transcript ids were retrieved using `biomaRt`^19^ and combined with canonical lists of TFs from UniProt (n= 1994) and FANTOM^67^ (n = 1988).

Groups were cross-referenced for uniqueness and annotated as following. TFs (n = 2662); coactivators were genes that exclusively associated with positive regulation of transcription (COA; n= 766); corepressors were genes that exclusively associated with negative regulation of transcription (COR; n = 599); mixed function coregulators were genes with evidence of context dependent negative and positive regulation of gene expression (MIXED; n = 511) (**Supplementary Table 1**).

### Testing family-wide changes in each gene category

Bootstrapping permutation approaches (boot), were used to test if the proportions of each gene group was altered more than predicted by chance. For mRNA and protein expression, Z-scores were calculated for gene expression data. In the cohorts of local tumor (MSKCC, TCGA) gene expression was calculated as Z-scores of tumor-normal comparisons with genes detectible in at least 80% of samples. In the other data sets the median expression data frame of RNA-Seq or Mass-Spec was converted to Z-scores. For copy number alterations (CNA) (via the GISTIC 2 method^53^) and mutations the data were used as obtained from cBioPortal^13^. To establish mutation frequency, the coding length of all exons for a given gene (including all alternative exons; BioMart) was calculated to yield the total gene CDS length. The mutation frequency was then calculated as the number of gene-length normalized mutations per gene and the square root of the summed mutation rate to control for the number of tumors in a cohort.

To test whether observed alteration frequencies within gene classes (e.g. COA, COR) was altered significantly, a vector of changes for all genes was calculated and the observed values (e.g. mean Z-score of expression) of a given class tested compared to all genes detected in each cancer cohort. Random sampling method was applied 100,000 times to select gene sets of equivalent size to simulate the distribution of changes across the genome for comparison^22^. Empirical p-values were calculated based on the group position relative to the sampling distribution of the genome.

### Testing relationships between the most significantly altered transcription factor and coregulators and clinical outcome

Gene expression levels in each gene family were filtered (`genefilter`) to select for genes that were commonly and significantly altered e.g. 2 Zscores in x % of tumors (figure legends). The expression and clustering of genes were then visualized with pheatmap, and the intersections visualized with `UpSetR`^17^ The association of patient cluster membership and clinical outcome (either categorical data or continuous data that was categorized) was then tested using a Chi-squared test and Kaplan-Meier curves generated for individual genes (`survival`) ^36^

### Analyses of the PGC1α-dependent transcriptome and cistrome in cells and TCGA cohorts

LNCaP_shPGC1A and LNCaP_shCtrl cells (1×10^6^ cells per well in a 6-well plate) adhered for 24 hr then dosed with ETYA (10 μM) or EtOH (vehicle control) and RNA extracted. All the samples were prepared in triplicates for RNA sequencing. Paired end sequence reads were aligned to the human genome (hg38) using `Rsubread`^46^, (> 90% of ~ 25×10^6^ unique reads/sample mapped) and translated to expression counts via `featurecounts`, followed by a standard edgeR pipeline^67^ to determine differentially expressed genes (DEGs), visualized with volcano plots (`ggplot2`) and interpreted by gene set enrichment^77^ and in particular the enrichment in the Hallmarks, Curated, GO and Reactome sets was analyzed.

Similarly, in the TCGA PRAD cohort tumors in the upper and lower quartile by PPARGC1A expression were identified and DEGs established. These DEGs were combined with publicly available PGC1α ChIP-Seq data^14^, and the enriched regions overlapped with PGC1α-dependent DEGs as indicated.

## RESULTS

### Transcription factor and coregulator groups are significantly down-regulated in localized prostate cancer cohorts

We generated unique gene lists of TFs (n = 2662); coactivators, (COA; n= 766); corepressors, (COR; n = 599); and mixed function coregulators, (MIXED; n = 511) (**Supplementary Table 1**). These groups were used to test the family-wide significance of changes in expression (mRNA or protein), CNA and mutation of each group in three local cancer (MSKCC, PRAD, OICR) and four advanced (MICH, FH, NEURO and SU2C) cancer cohorts (**Table 1**).

Hypergeometric tests revealed that COAS, CORS and MIXED genes, but not TFs, were significantly protected (p < 0.01) from loss of function structural variation in normal tissues (GNOMAD^70^) and significantly enriched in Pan-Cancer fitness genes^5^.

Next, a bootstrapping permutation approach was used to test if observed family-wide changes were more than predicted by chance ^49^. We calculated empirical p-values based on the position of the family gene set (e.g. TF, COA etc) relative to the distribution of random gene sets of the same size. The heatmap in **Supplementary Figure 1A** shows CORS expression in the TCGA cohort, indicating the genes are commonly down-regulated. To test CORS down-regulation we identified the proportion of these genes altered by > 2 Z-scores (vertical dashed red line, **Supplementary Figure 1A, right panel**) and compared this to the proportion of all gene families down-regulated in sets of the same size (green histogram); a similar analyses was applied to the up-regulated genes (red histogram). The observed proportion of CORS down-regulated by > 2 Z-scores was significantly more (and hence, to the right) than the estimated proportion of down-regulated groups of the same size and randomly sampled (p=1e^-05^). The CNA data and normalized exome mutation rate were treated in the same manner. Positive controls included gene groups known to be significantly altered by expression (nuclear hormone receptor (NR) down-regulation; HOX family (HOX)^49^ up-regulation), mutation (Cosmic_mutant) (**Supplementary Figure 1B**) or CNA (COSMIC_CNA) (data not shown) ^80^.

To visualize significant results we plotted the negative log_10_ of the FDR-corrected empirical pvalues for each test across cohorts (**Figure 1**). The mRNA expression in MSKCC and TCGA, and protein expression in OICR were significantly more down-regulated and/or less up-regulated. For example, CORS were significantly more down-regulated (MSKCC, TCGA, FH) and less up-regulated (MSKCC, FH, OICR) than predicted by chance. Indeed, all groups were more down-regulated or less-upregulated in at least 4 cohorts. Only two group/cohort tests revealed significant up-regulation; COR genes in MICH, and MIXED genes in OICR.

**Figure 1.**
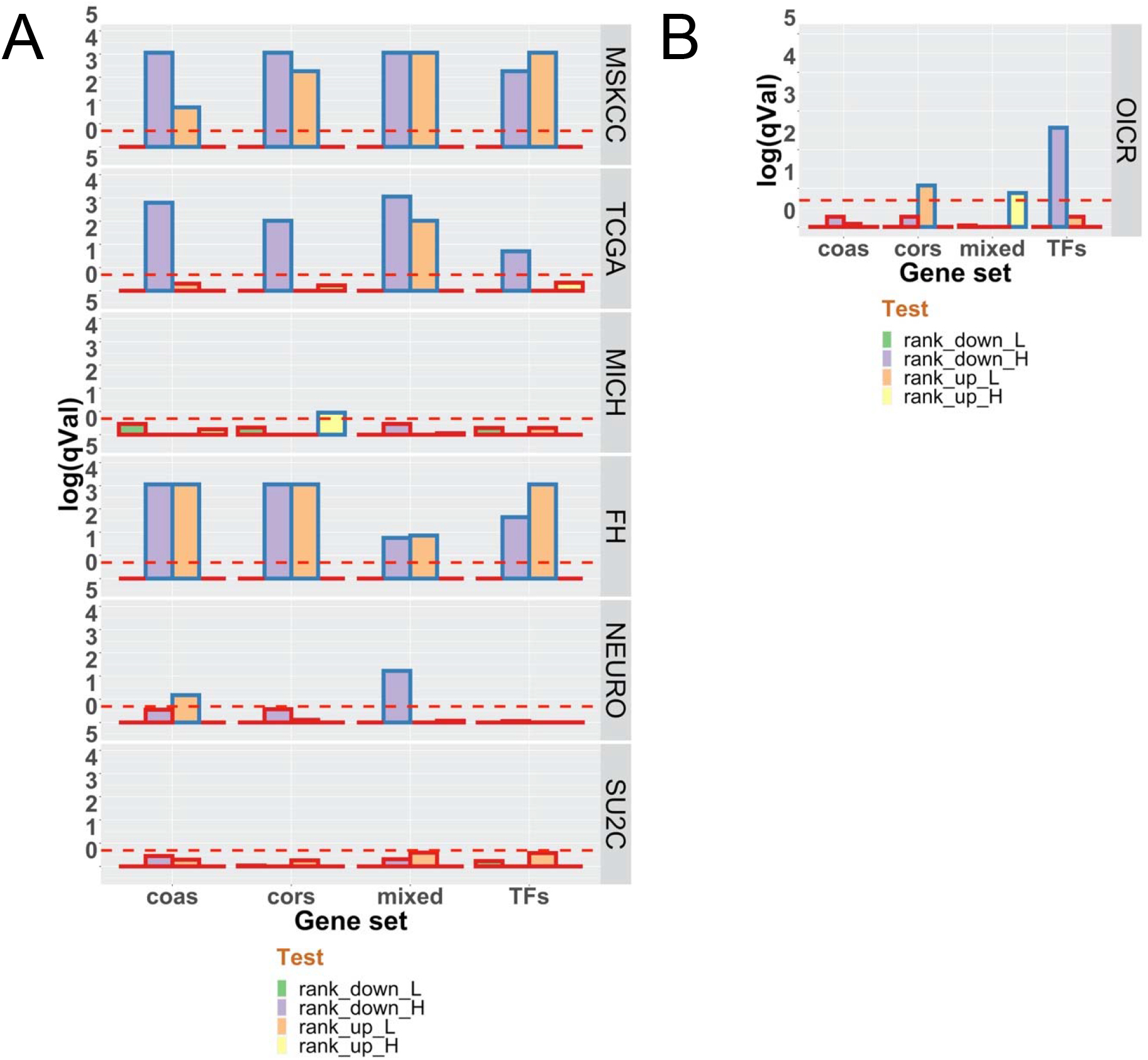
Family-wide analyses of transcription factors (TFs), coactivators (COAS), corepressors (CORS) and mixed function coregulators (MIXED) in PCa cohorts. The mean rank of the genes in each family were compared to the distribution of ranking of the same number of genes randomly sampled 100,000 times. The empirical p-values were −log10 transformed and FDR corrected for mRNA expression (**A**) and proteins (**B**).

Therefore down-regulation of these groups was more common than predicted by chance in local tumors. The OICR cohort displayed significant protein down-regulation of TFs, although the protein detection of all genes is significantly reduced in number (**Table 1**) and therefore caution is needed to compare between patterns of RNA and protein expression. Nonetheless, the correlation between RNA (TCGA or SU2C) and protein (OICR) when considering RNA transcripts for detected proteins ranged from ρ 0.4 to 0.55 (**Supplementary Figure 2**).

**Table 1:** Summary of the prostate cancer cohorts. The number of tumors and the number for genes detected from each category in the unique gene lists (TFs (n = 2662); coactivator (COA; n= 766); corepressor (COR; n = 599); mixed function coregulator (MIXED; n = 511).

| Cohort | TumorType | Material | Tumors | COAS | CORS | MIXED | TFs |
| --- | --- | --- | --- | --- | --- | --- | --- |
| MSKCC | Local | RNA, CNA | 152 | 696 | 439 | 457 | 1506 |
| TCGA | Local | RNA, CNA, Exome muts | 498 | 663 | 415 | 414 | 1305 |
| OICR | Local | Protein | 76 | 320 | 185 | 156 | 295 |
| FH | Metastatic | RNA, CNA, Exome muts | 148 | 712 | 445 | 465 | 1384 |
| MICH | Metastatic | RNA, CNA, Exome muts | 95 | 276 | 179 | 169 | 740 |
| SU2C | Metastatic | RNA, CNA, Exome muts | 271 | 543 | 343 | 384 | 1122 |
| NEURO | Metastatic | RNA, CNA, Exome muts | 51 | 725 | 466 | 485 | 1518 |

By contrast to the findings on RNA and protein expression, the mutation and CNA data were largely negative; COAS and MIXED genes were more mutated than predicted by chance in the FH cohort only (**Supplementary Figure 1C**). These findings suggest that changing the stoichiometry of TF and coregulator interactions is potentially more impactful in local tumors than disruption through either mutation or CNA.

### Reduced expression of transcription factors and coregulators associates with more aggressive PCa

Next, from each of the 15 significant RNA expression-cohort relationships we identified those genes with the most frequent and greatest expression change. For example, COAS in the TCGA cohort, were filtered for genes altered by > 2 Z-scores in 35% of TCGA tumors. The filtered RNA (**Figure 2**) and protein expression (**Supplementary Figure 3**) were visualized as heatmaps.

**Figure 2.**
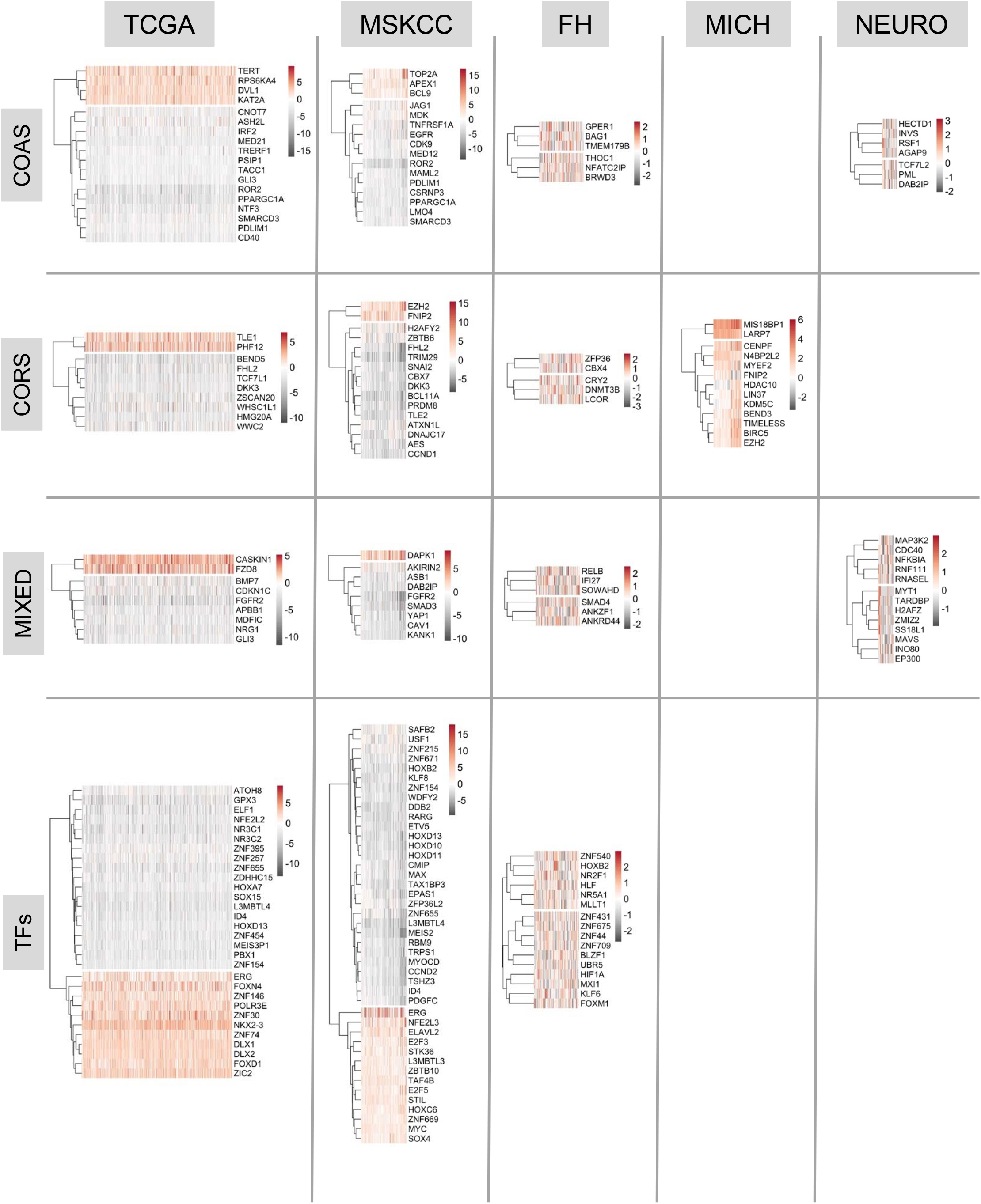
Expression of transcription factors (TFs), coactivators (COAS), corepressors (CORS) and mixed function coregulators (MIXED) in PCa cohorts. Genes in each group were filtered (`genefilter`; >2.5 Z scores in 35% tumors) to reveal the most frequently and strongly altered and genes and tumors (columns) were visualized (`pheatmap`).

Shared targets between the TCGA and MSKCC cohorts were investigated further (**Figure 3A**). PubMed analyses (search terms given in Table Legend) revealed an uneven representation of these genes in the context of PCa (**Table 2**). These genes included those known to be commonly altered, including *ERG*, a frequently up-regulated a TF in PCa^61^ and common target of translocations with TMPRRS2^69, 85^. By contrast, to date, the COA, PDLIM1 has not been investigated in the context of PCa, and others such as *PPARGC1A* have only been modestly investigated^20, 38, 73, 83, 88, 90, 98^.

**Figure 3.**
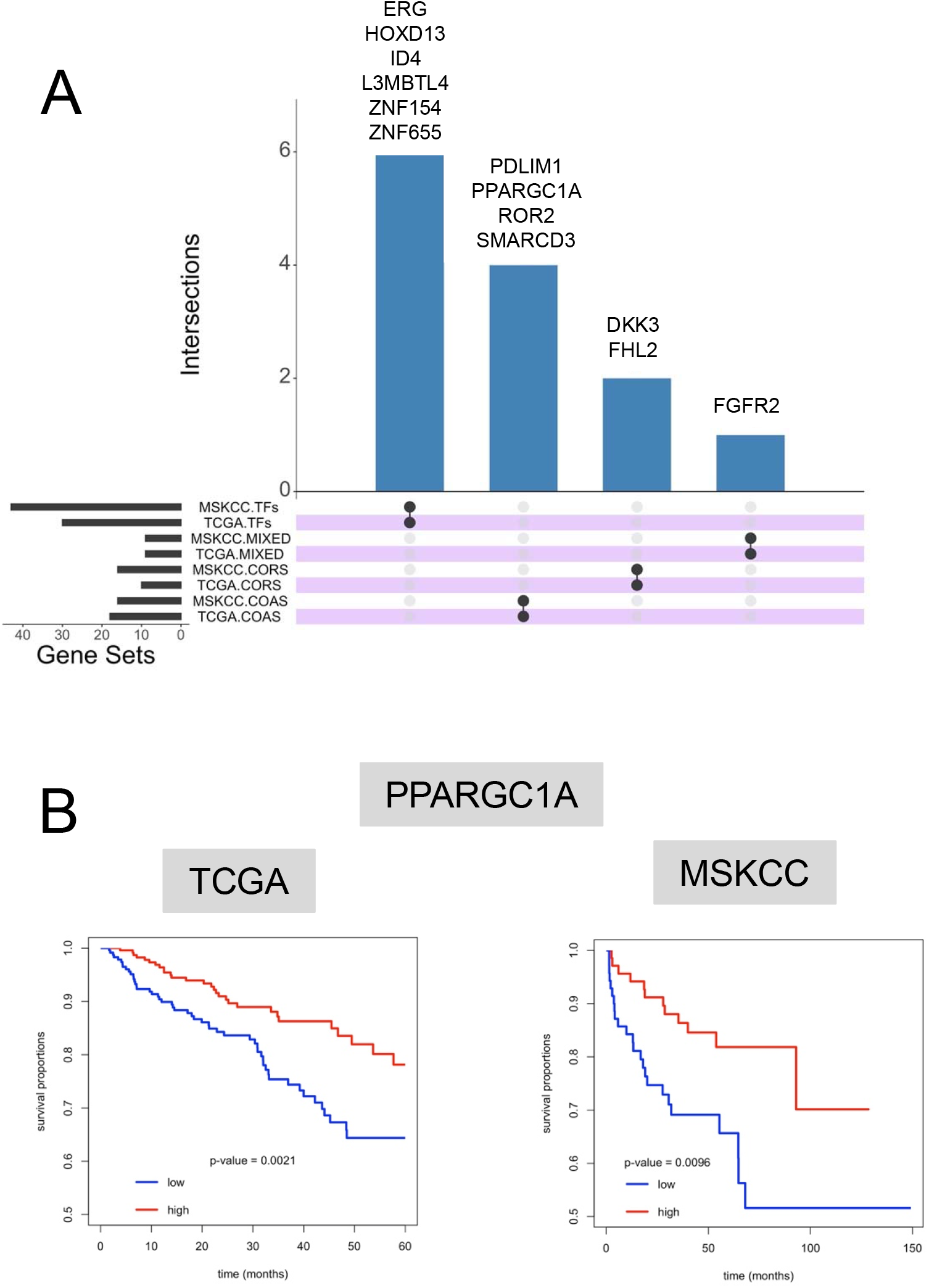
Identification of commonly altered transcription factors (TFs), coactivators (COAS), corepressors (CORS) and mixed function coregulators (MIXED) in the MSKCC and TCGA cohorts. **A.** Genes commonly altered in both cohorts were identified and visualized (UpsetR). **B.** Kaplan-Meier plots of the relationship between tumors with lower and upper quartile PPARGC1A expression and time to biochemical progression.

**Table 2:** Number of publications addressing the top altered genes in both the TCGA and MSKCC cohorts. The genes were mined in PubMed using the following search term (prostate cancer[Title/Abstract]) AND *gene name*[Title/Abstract])

| SharedGenes | Group | Publications |
| --- | --- | --- |
| ERG | TFs | 1057 |
| HOXD13 | TFs | 20 |
| ID4 | TFs | 0 |
| L3MBTL4 | TFs | 3 |
| ZNF154 | TFs | 0 |
| ZNF655 | TFs | 0 |
| PDLIM1 | COA | 0 |
| PPARGC1A | COA | 7 |
| ROR2 | COA | 4 |
| SMARCD3 | COA | 0 |
| DKK3 | COR | 13 |
| FHL2 | COR | 16 |
| FGFR2 | MIXED | 45 |

Next, we examined how expression related to PCa progression by generating Kaplan Meier estimates of the time to biochemical progression (**Table 3**). In both TCGA and MSKCC cohorts, *PPARGC1A* was the only gene to be commonly and significantly down-regulated, and significantly associated with reduced time to biochemical progression (**Figure 3B**).

**Table 3:**
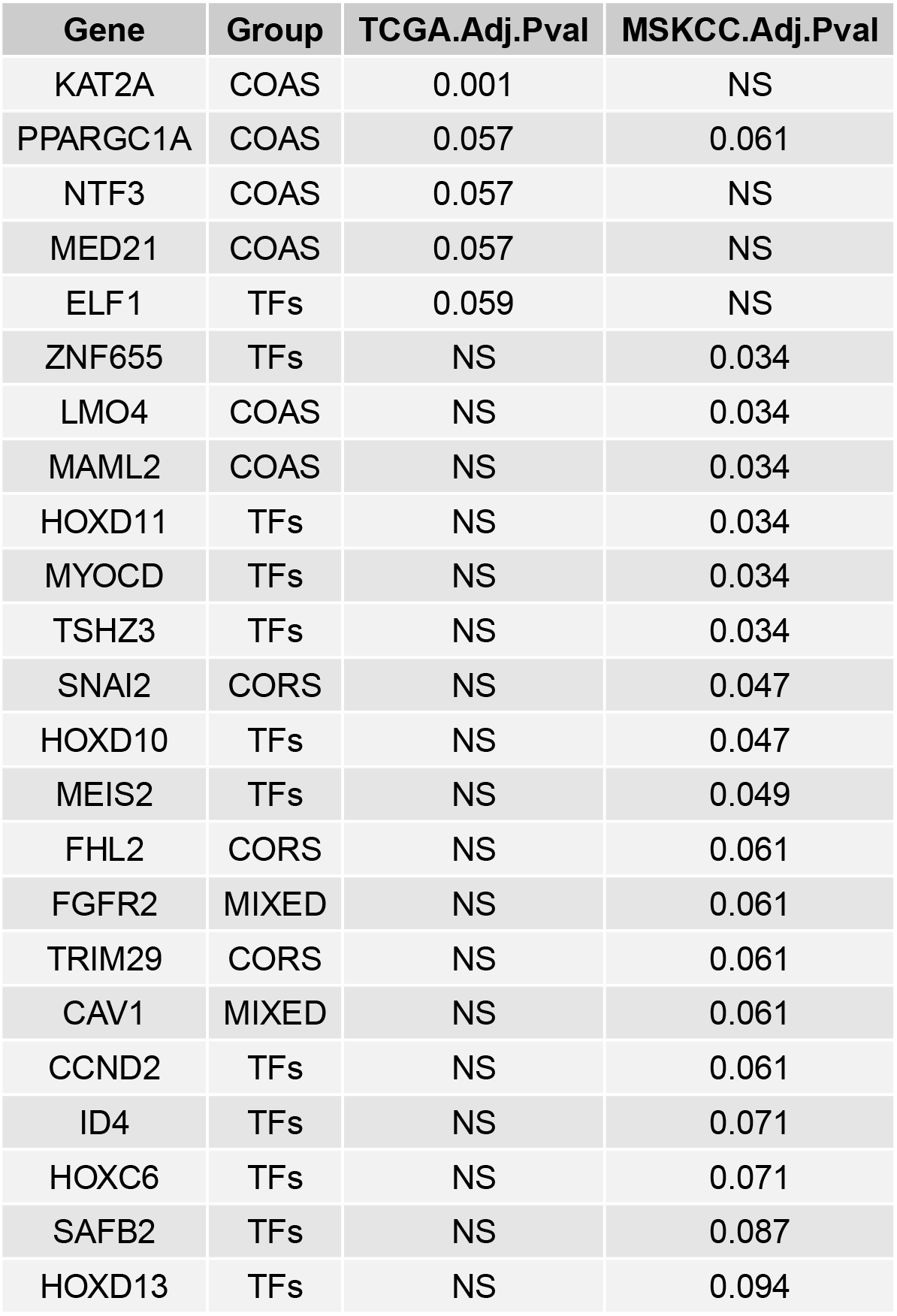
Relationships between altered expression and time to disease progression. Kaplan-Meier plots of the relationship between tumors with lower and upper quartile expression of the indicated gene and time to biochemical progression were generated and the FDR-corrected pvalues indicated.

| Gene | Group | TCGA.Adj.Pval | MSKCC.Adj.Pval |
| --- | --- | --- | --- |
| KAT2A | COAS | 0.001 | NS |
| PPARGC1A | COAS | 0.057 | 0.061 |
| NTF3 | COAS | 0.057 | NS |
| MED21 | COAS | 0.057 | NS |
| ELF1 | TFs | 0.059 | NS |
| ZNF655 | TFs | NS | 0.034 |
| LMO4 | COAS | NS | 0.034 |
| MAML2 | COAS | NS | 0.034 |
| HOXD11 | TFs | NS | 0.034 |
| MYOCD | TFs | NS | 0.034 |
| TSHZ3 | TFs | NS | 0.034 |
| SNAI2 | CORS | NS | 0.047 |
| HOXD10 | TFs | NS | 0.047 |
| MEIS2 | TFs | NS | 0.049 |
| FHL2 | CORS | NS | 0.061 |
| FGFR2 | MIXED | NS | 0.061 |
| TRIM29 | CORS | NS | 0.061 |
| CAV1 | MIXED | NS | 0.061 |
| CCND2 | TFs | NS | 0.061 |
| ID4 | TFs | NS | 0.071 |
| HOXC6 | TFs | NS | 0.071 |
| SAFB2 | TFs | NS | 0.087 |
| HOXD13 | TFs | NS | 0.094 |

### Knockdown of PGC1α increases proliferation and alters a large transcriptome

We generated stable PGC1α knockdown clones in LNCaP cells and examined the cell phenotype, and basal and PPARγ ligand ETYA transcriptome^4, 91^. (**Figure 4A**). Reduced PGC1α expression increased the basal cell proliferation rate compared to vector controls, but not the anti-proliferative effect of ETYA (**Figure 4B**). Reduced PGC1α also increased the invasiveness of the cells as measured by a scratch assay (**Figure 4C**).

**Figure 4.**
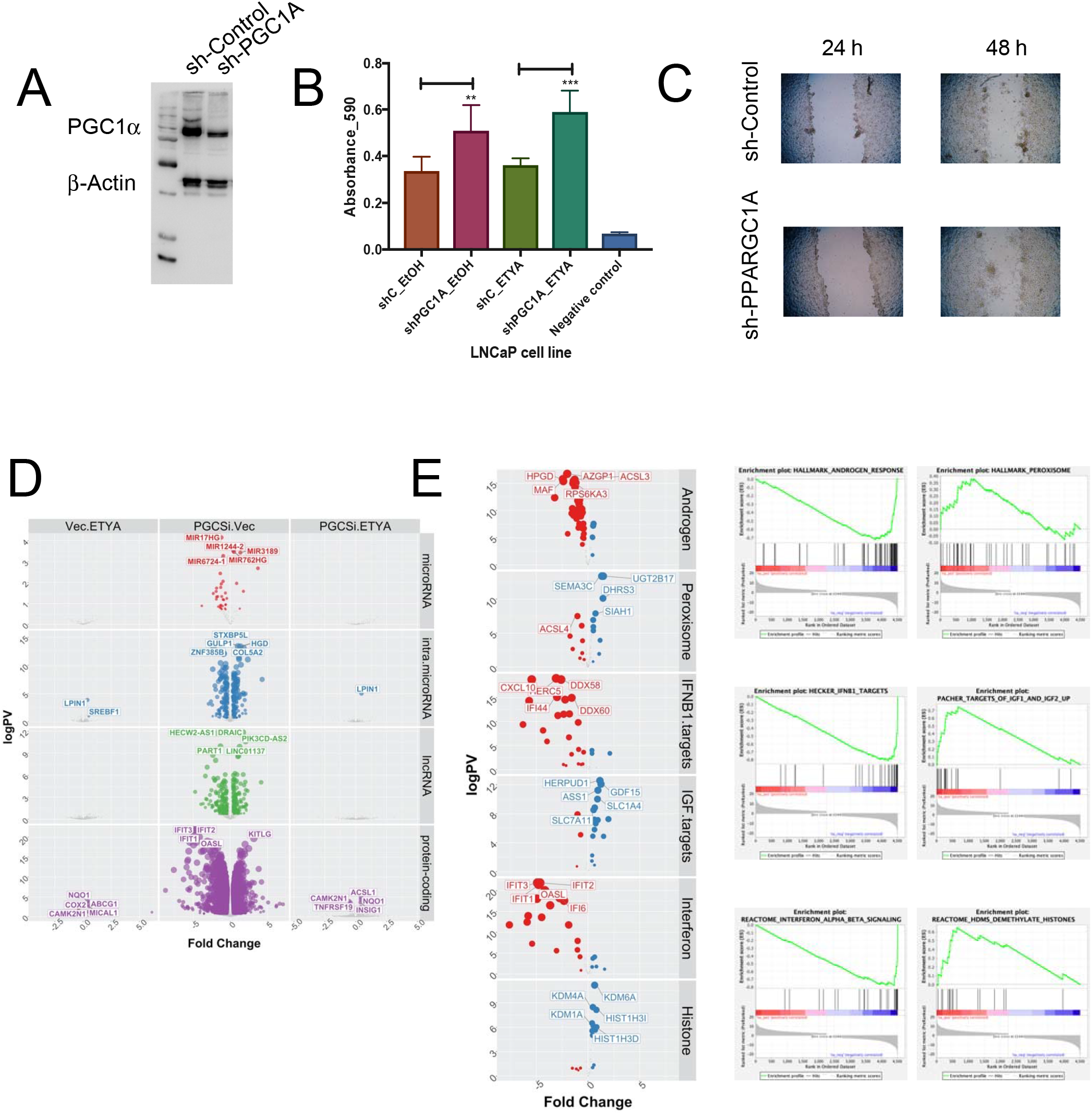
Stable knockdown of PGC1α in LNCaP cells changes proliferation and gene expression patterns. **A.** LNCaP cells were each stably transfected with two shRNA constructs (sh-PGC1A 34 and 35) targeting *PPARGC1A* resulting in reduced PGC1α expression (sh-PGC1A) compared to empty vector (sh-CTRL) at protein level as detected in western blotting. **B**. Measurements of cellular levels of ATP, as an indicator of cell viability was detected in the vector controls and knockdown cells. Each measurement was performed in biological triplicates and in triplicate wells. Cells were treated in triplicate with exogenous PGC1α ligand ETYA (10 μM, 96hr) or EtOH vehicle control. Increased cell proliferation was seen in sh_PGC1A cells after treatment with ETYA at 96 hr. **C**. Time course scratch closure of sh-Control and sh-PPARGC1A cells mechanically wounded with p200 sterile pipette tip, sh-PPARGC1A after 48 hr showed increased cell migration compared to sh-Control. **D.** LNCaP sh-PGC1A and sh-CTRL cells were treated with ETYA (10 μM, 24hr) or EtOH vehicle control and total RNA expression and RNA-Seq undertaken according to the `edgeR` pipeline. Volcano plots depicting expression changes upon PGC1α knockdown or in response ETYA in the indicated classes of RNA in color (-log10(p.adj) >1, abs (log2(fold change))). **E**. Summary of significantly enriched pathways from gene set enrichment analyses (GSEA) (FDR q.val < 0.05) associated with reducing PGC1α expression levels.

RNA-Seq revealed the impact of reduced PGC1α expression in LNCaP cells had a profound impact on gene expression, reflecting the significant impact on proliferation the knockdown of PGC1α (**Figure 4D, E**). These changes included 37 microRNA, 243 intra.microRNA, 343 lncRNA and 3600 protein coding genes significantly altered. By contrast the impact of ETYA was very modest. This suggests that PGC1α regulation of ligand-activated PPARγ does not appear to be significant, at least with respect to the impact of ETYA exposure (**Figure 4D**).

Supporting a coactivator function, PGC1α knockdown down-regulated more than up-regulated genes. For example, 30 miRNA were down-regulated (e.g. miR17 host gene) and only 7 up-regulated, and similarly 1993 protein-coding genes were down-regulated and 1607 up-regulated. The fold change was also skewed to down-regulation, with the mean logFC for down-regulated protein-coding genes being −1.11, and 0.83 for up-regulated genes and similarly the mean logFC for down-regulated lncRNA was −1.20, and 0.87 for up-regulated genes (**Figure 4D**).

We undertook GSEA analyses to classify the PGC1α-dependent changes in gene expression ^76^. **Supplementary Figure 4** illustrates the top eight positive and negative significant (FDR < .1) normalized enrichment scores (NES) in four gene set categories; Hallmarks, Curated, GO and Reactome sets. These highly enriched gene patterns illuminate the cancer biology impact of reduced PGC1α expression. The NES plots and expression of the altered genes in each term is shown in **Figure 4E**.

Analyses of the Hallmark gene sets revealed androgen response genes were negatively and peroxisome genes positively enriched, respectively, suggesting that PGC1α expression represses AR signaling and enhances the activity of the peroxisome. AR target genes notably associated with energy production were repressed, including the putative tumor suppressor hydroxyprostaglandin dehydrogenase (*HPGD*)^29^, as well as Acyl-CoA synthetase long chain family member 3 (ACSL3)^55^ and Alpha-2-glycoprotein 1, zinc-binding (*AZGP1*)^12^. Up-regulated peroxisome genes included retinoic acid anabolizing enzyme, Dehydrogenase/Reductase 3 (*DHRS3*)^48^, and the steroid catabolizing enzyme UDP Glucuronosyltransferase Family 2 Member B17 (*UGT2B17*)^99^. These changes in hallmark gene sets suggests that altered PGC1α disrupts the energetic utilization of the cell and distorts AR signaling.

This was also borne out by changes in Curated gene sets, with repression of Interferon signaling genes including retinoic acid inducible DExD/H-Box Helicase 58 (*DDX58*)^16^, and upregulation of IGF-signaling gene targets including the AR-target genes Homocysteine Inducible ER Protein With Ubiquitin Like Domain 1 (*HERPUD1*)^71^ and Growth Differentiation Factor 15 (*GDF15*), known to be a prostate mitogen^94^. Similarly, Reactome gene set enrichment revealed repression of Interferon regulated genes including Interferon Induced Protein With Tetratricopeptide Repeats 1 (*IFIT1*)^62^, which is actually repressed by PPARα^31^. Interestingly, this was accompanied by upregulation of a number of histone modifying enzymes including lysine demethylases^31^ (*KDM1A, KDM4A, KDM6A*) suggesting a potential for significant changes in the methylation status of histone lysine modifications such as H3K4me3 and H3K27me3, and thereby impacting the epigenome.

### Genes bound and regulated by PGC1α associate with more aggressive PCa

Reduced PGC1α expression impacted gene expression positively and negatively. Whether PGC1α exerts both coactivator and corepressor functions is obscured by direct (*cis*) and indirect (*trans*) relationships between PGC1α and the regulated genes captured by RNA-Seq. To identify PGC1α *cis*-dependent gene expression we examined how genes bound by PGC1α binding relate to changes in gene expression.

PGC1α ChIP-Seq data^14^ were combined with the PGC1α-dependent gene expression patterns and shared genes were identified. Given each protein-coding genes is regulated by multiple enhancers^65, 84^ and promoter-capture Hi-C experiments revealed the median distance between enhancer and target gene is 158 kb^32^. Therefore PGC1α ChIP-Seq peaks were segregated into different chromatin states identified in LNCaP cells (e.g. Promoter, Active Enhancer)^78^, and annotated to known genes within 100 kb. Thus, the 1304 PGC1α ChIP-Seq peaks overlapped with 914 chromatin states identified in LNCaP, and annotated to 2381 peak:gene relationships.

In the first instance we examined how *PPARGC1A* correlated either to genes that were differentially regulated by PGC1α knockdown in LNCaP cells (n=4187, **Figure 4D**) and bound by PGC1α overlapping with LNCaP chromatin states (n=2381). We compared the relationships between PGC1α binding and gene expression in the TCGA cohort. In the first instance cumulative correlation plots were generated, which revealed that the empirical correlation between the expression of *PPARGC1A* and genes regulated by *PPARGC1A* shRNA were significantly more positive than the background of all expressed genes (KS test; p<1e-9) (**Supplementary Figure 5**, green symbol). This suggests that there is a stronger positive correlation between PGC1α expression and PGC1α-dependent genes than predicted by chance, and fits with a model of coactivator function. By contrast, the correlation between the expression of *PPARGC1A* and genes directly bound by PGC1α were significantly more negative than predicted by chance and suggested the correlations were significantly more negative (KS test; p<1e-9). This suggests that direct binding of PGC1α is more nuanced in its direct relationship with gene levels, with a more even distribution of negative and positive correlations with expression.

Next, these 2381 PGC1α peak:gene relationships linked to LNCaP chromatin states were filtered to include genes that were: 1. differentially regulated by *PPARGC1A* shRNA in LNCaP cells (n = 4187, **Figure 4**); 2. differentially expressed in the TCGA cohort between the top and bottom quartile expressing *PPARGC1A* expression (n = 7324). This identified 160 genes bound by PGC1α and significantly altered by PGC1α knockdown in LNCaP cells and significantly altered *PPARGC1A*-dependent expression in TCGA, with the most altered 63 genes in the TCGA cohort shown in **Figure 5A**. Expression genes clustered higher Gleason grade tumors (X-squared = 5.0601, p-value = 0.025).

**Figure 5.**
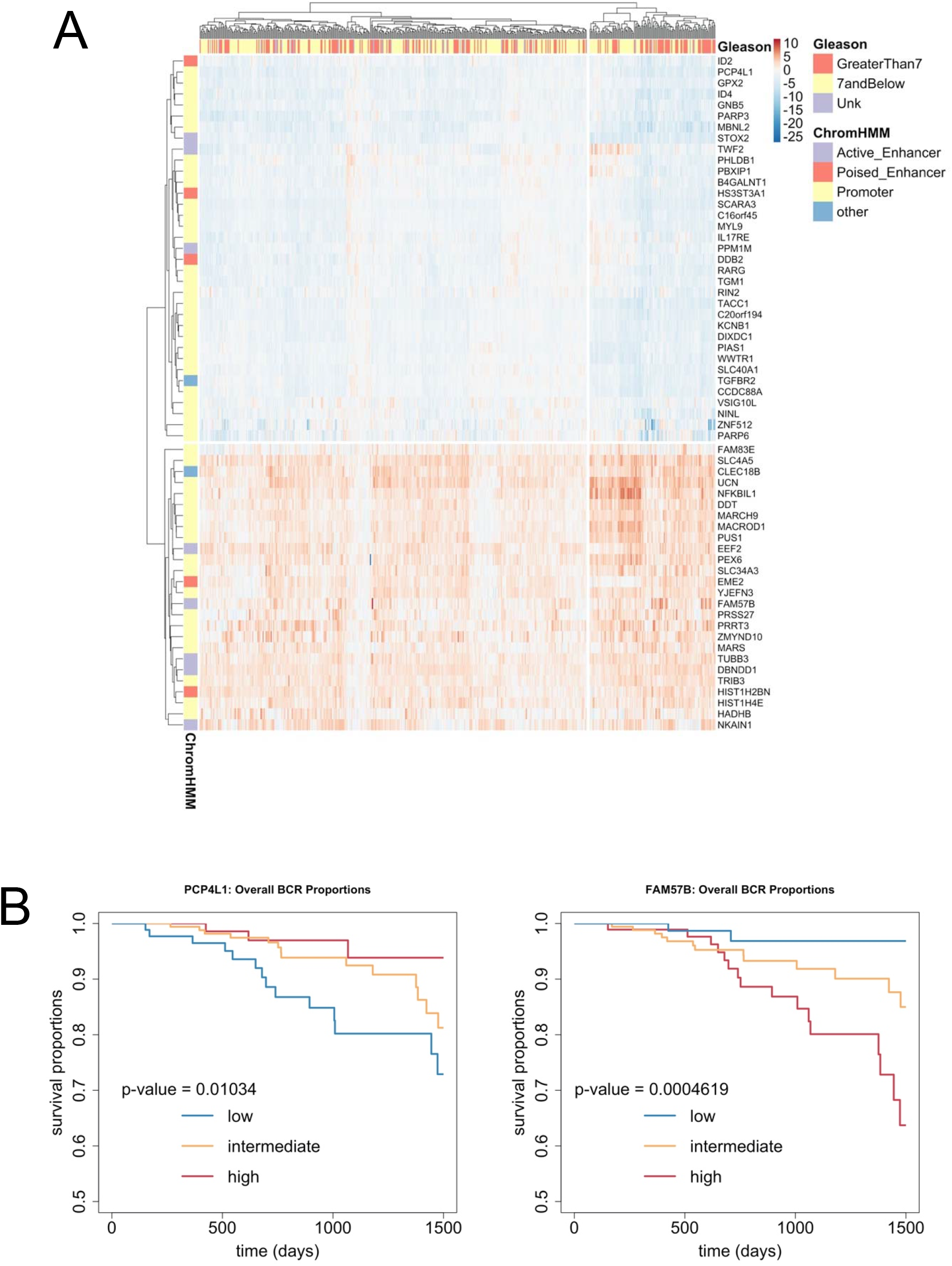
Expression of PGC1α-regulated and PGC1α-bound genes in the TCGA cohort. **A.** PGC1α ChIP-Seq data (GSE75193) were annotated to genes within 100 kb, to yield PGC1α peak:gene relationships. These were then filtered to include only those that overlapped by 1 bp with chromatin states in LNCaP derived using ChromHMM to yield PGC1α peak:gene relationships. Thus, the 1304 PGC1α ChIP-Seq peaks overlapped with 914 specific LNCaP chromatin states and annotated to 2381 peak:gene relationships. In turn these further filtered to include only those genes that were: 1. differentially regulated by PPARGC1A shRNA in LNCaP cells (n = 4187, **Figure 4**); 2. differentially expressed in the TCGA cohort between the top and bottom quartile expressing PPARGC1A expression (n = 7324), resulting in 160 genes that were significantly bound by PGC1α and whose expression was significantly altered by both shRNA PPARGC1A in LNCaP cells and significantly altered in TCGA tumors in PPARGC1A-dependent manner. The top 63 genes are illustrated (`pheatmap`) and their expression patterns clustered higher Gleason grade tumors (X-squared = 5.0601, p-value = 0.025). **B.** Kaplan–Meier plots of the relationship between tumors with lower and upper quartile expression of the indicated genes and time to biochemical progression.

Interestingly, amongst the PGC1α-dependent gene signature 25 genes were upregulated, and 38 genes were actually downregulated, again suggesting that the direct impact of PGC1α on gene expression is more nuanced and not exclusively a coactivator. Finally, we screened how expression of these 63 PGC1α-dependent genes related to tumor outcome by generating Kaplan–Meier plots. The top five significant genes were *FAM57B, EME2, DDB2, PCP4L1* and *PBXIP1* (**Supplementary Table 2**) and the plots for an upregulated gene (*FAM57B*) and a downregulated gene (*PCP4L1*) are shown in **Figure 5B**. Interestingly, *FAM57B* is an established PPARγ target gene that regulates adipocyte differentiation^97^. Similarly, *PCP4L1* is related to obesity induced phenotypes^37^ and *PBXIP1* plays a role in regulation of differentiation^51^ in part by regulating ERα signaling^95^. By contrast *EME2* and *DDB2* relate to DNA repair^52, 82^.

## DISCUSSION

The current study was undertaken to identify the most significantly altered and clinically-relevant TF and co-regulators in PCa. It was reasoned that this knowledge may help to explain disease progression risks, or to highlight novel therapeutic opportunities. Assessment of family-wide significance across seven PCa cohorts revealed that TFs, COAS, CORs and MIXED were more down-regulated and less up-regulated at the mRNA and protein level, most clearly in local tumor cohorts. In contrast, family-wide CNA and mutation levels were not significant. Filtering those group/cohort relationships identified the most altered genes in the local tumor cohorts (MSKCC and TCGA). Overlapping identified commonly altered genes associated with PCa including *ERG*^61^ (which was also altered at the protein level in the OICR cohort) and *FGFR2*^44^, but also identified others that were relatively under-investigated such as *PPARGC1A*^11, 88, 89^ but associated with worse disease free survival, or uninvestigated in PCa, such as the metastasis suppressor *PDLIM1*^30^. Stable knockdown of PGC1α increased proliferation and invasion of LNCaP cells, and profoundly altered the transcriptome. Finally, PGC1α bound and regulated genes associated with higher grade tumors in the TCGA cohort and individual genes such as the PPARγ target gene *FAM57B* associated with worse disease-free survival.

The global down-regulation of TFs and co-regulators suggests that an initial event in PCa is disrupting the flexibility of gene regulatory signaling that may limit the permutations of TF co-regulator interactions or lessen the ability of the TF and co-regulators to form effective stoichiometric interactions for correct functioning. This may suggest that although these genes are reduced in expression they are not altered by structural variation, and therefore they remain functional, and in turn this may support the disrupted stoichiometry argument. The current findings would suggest that globally the flexibility of TF actions is disrupted. This may suggest that signaling is more restricted in terms of capacity, and this is seen in the PCa AR transcriptome ^27,29^, but also in signaling via MYC and AP1 (reviewed in^73^).

The stringent filtering of these data and overlap between TCGA and MSKCC revealed commonly and significantly altered genes but are unevenly investigated in PCa (**Table 2**). Genes that are only identified in one cohort and whose expression is associated with more aggressive disease (**Table 3**) reveals a number of genes that are potentially impactful in disease but are under-investigated or have not been investigated at all in the context of PCa. *PPARGC1A* was selected for mechanistic study, given it was down-regulated and significantly associated with worse disease-free survival in both MSKCC and TCGA cohorts. Knockdown of PGC1α in LNCaP cells increased proliferation and migration, coupled with a profound impact on the transcriptome. The PGC1α transcriptome was enriched for terms associated with AR signaling as well overlaps with regulation of the peroxisome, interferon signaling and epigenetic modifiers. These findings were further supported when considering target genes that were bound by PGC1α and differentially expressed in the TCGA cohort. These genes distinguished aggressive tumors and included RARγ/TACC1^50^, which we have previously established to act to antagonize AR signaling, as well as genes associated PPARγ signaling. Furthermore, motif analyses revealed that basic helix-loop-helix (b-HLH) motifs were enriched in the PGC1α-ChromHMM sites, as were sites for LXR binding. These findings suggest that PGC1α regulates signaling by a cohort of nuclear receptors including AR, RARγ, PPARγ as well epigenetic process and interferon signaling.

Earlier studies by Carracedo and colleagues identified reduced expression of PGC1α in PCa, and these workers pursued a strategy to over-express PGC1α in advanced PCa cell (PC-3) cells line and analyzed these effects in two impactful studies^88, 89^. Their findings support a role for PGC1α to regulate a transcriptome that is regulated by PGC1α and ERRα cross-talk, which regulates MYC signaling to regulate invasiveness; indeed in the current study b-HLH motifs were identified in the PGC1α-binding sites identified in LNCaP cells. The PGC1α-dependent transcriptome in PC-3 cells is ~ 900 genes (GSE75193) and few (n=119) appear to be shared with the RNA-Seq data generated in the current study, but none of the genes proposed by the authors as a PGC1α-dependent signature overlap with the 63 PGC1α-dependent and PGC1α-bound genes significantly altered in TCGA cohort associated with more aggressive disease.

Thus, whilst the current study and the Carracedo studies both identify important roles for PGC1α to regulate tumor aggressiveness, the mechanisms appear to be different. This may arise for technical reasons (for example, cell background and transcriptomic approach) and may reflect the differential impact of knockdown versus over-expression selecting for different transcriptionaly-dependent events. Certainly, the fact that PGC1α-dependent and PGC1α-bound genes favor down-regulated over up-regulated genes (~ 2:1, **Supplementary Table 3**) suggests that PGC1α participates in diverse transcriptional events. Outside of PCa, in renal disease it has been proposed that PGC1α determines phenotypic consequences by selecting which TFs to interact with, and that RARs/RXRs, PPARs is associated with fatty acid metabolism^45^. It is tempting to speculate the phenotype is current study is associated with similar energetic changes.

A final comment concerns the scale of the analyses and the relatively generic approach. This approach considers all genes in a class or superfamily to identify which may have merit to investigate. This is an important step for many wet-lab approaches, given that it can often take considerable resources to dissect a gene function. For example, the 20,000 ft view approach to cancer genomics in PCa integrates genome-wide data to identify novel networks ^18,58^. Traditionally trained wet-lab based investigators often face challenges in assimilating these findings and may rather search for gene(s) in the supplementary data that are from the family on which they study; this can be considered as a 200 ft view. The current approach is neither the 200 ft nor 20,000 ft views, but rather a 2,000 ft view. That is, the approach allows an investigator to remain focused in their research arena, for example TF co-regulators, but ask the global question over how these are altered and at what stage of cancer progression. This is a relatively generic approach and may find traction with investigators across disease types.

## Acknowledgements

*MJC* acknowledges support in part from the Prostate program of the Department of Defense Congressionally Directed Medical Research Programs [W81XWH-14-1-0608], the National Institute of Health Cancer Center Support Grant (P30CA016058) to the OSUCCC The James. *EW* and *MJC* acknowledge seed-funding support from the Molecular Carcinogenesis program of the OSUCCC The James, CCSG. *CLB, MJC, DAL* acknowledge support in part from Prostate Cancer, UK [RIA18-ST2-022]. *EAM* acknowledges support The Ohio State University Translational Data Analytics Institute, startup funds from The Ohio State University.

