## Supplementary material for "Genome-wide analyses of transcription factors and co-regulators across seven cohorts identified reduced PPARGC1A expression as a driver of prostate cancer progression": Supp Figures

### Slide 1
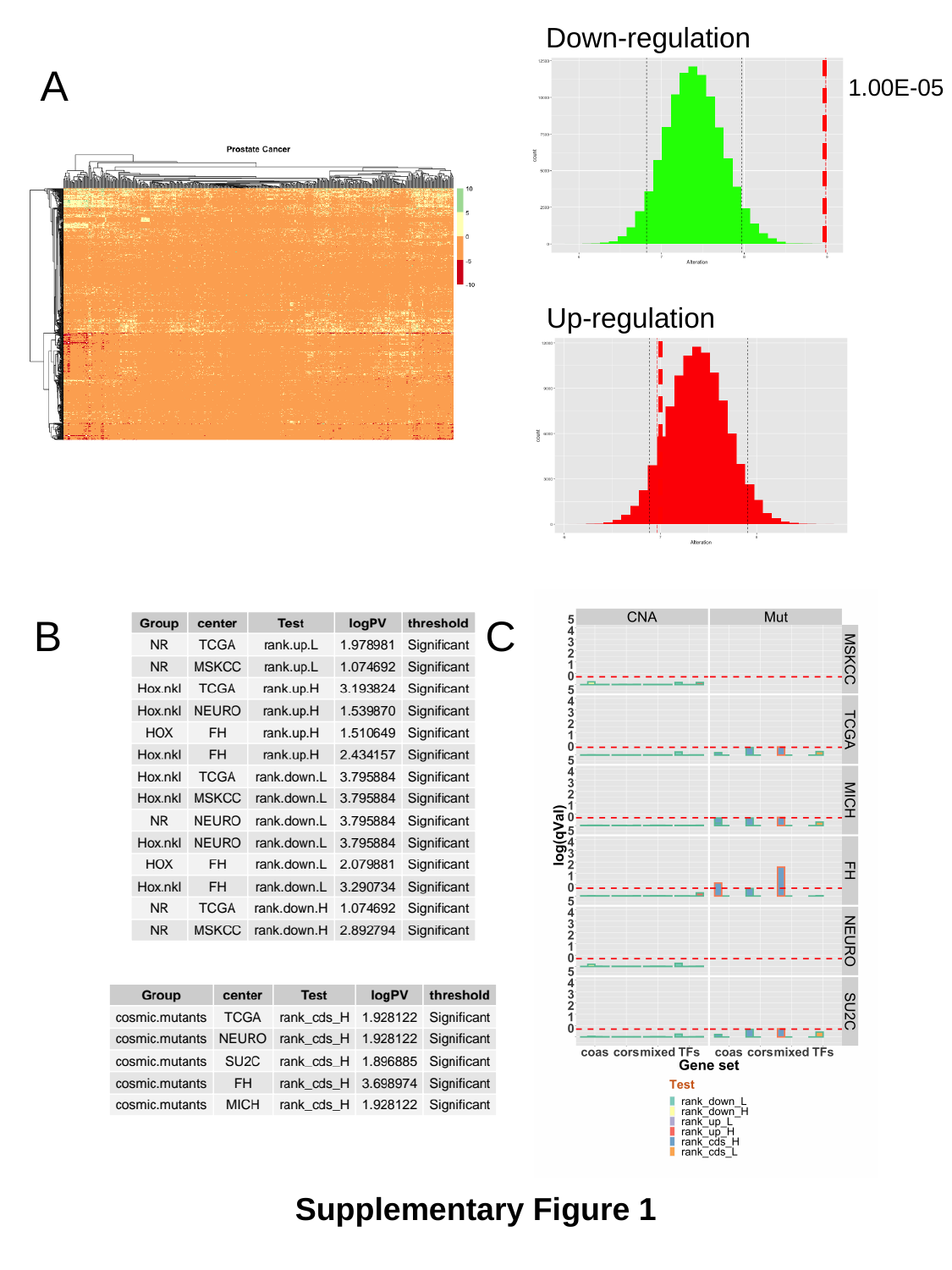

Down-regulation
1.00E-05
Up-regulation
A
C
B
Supplementary Figure 1

### Slide 2
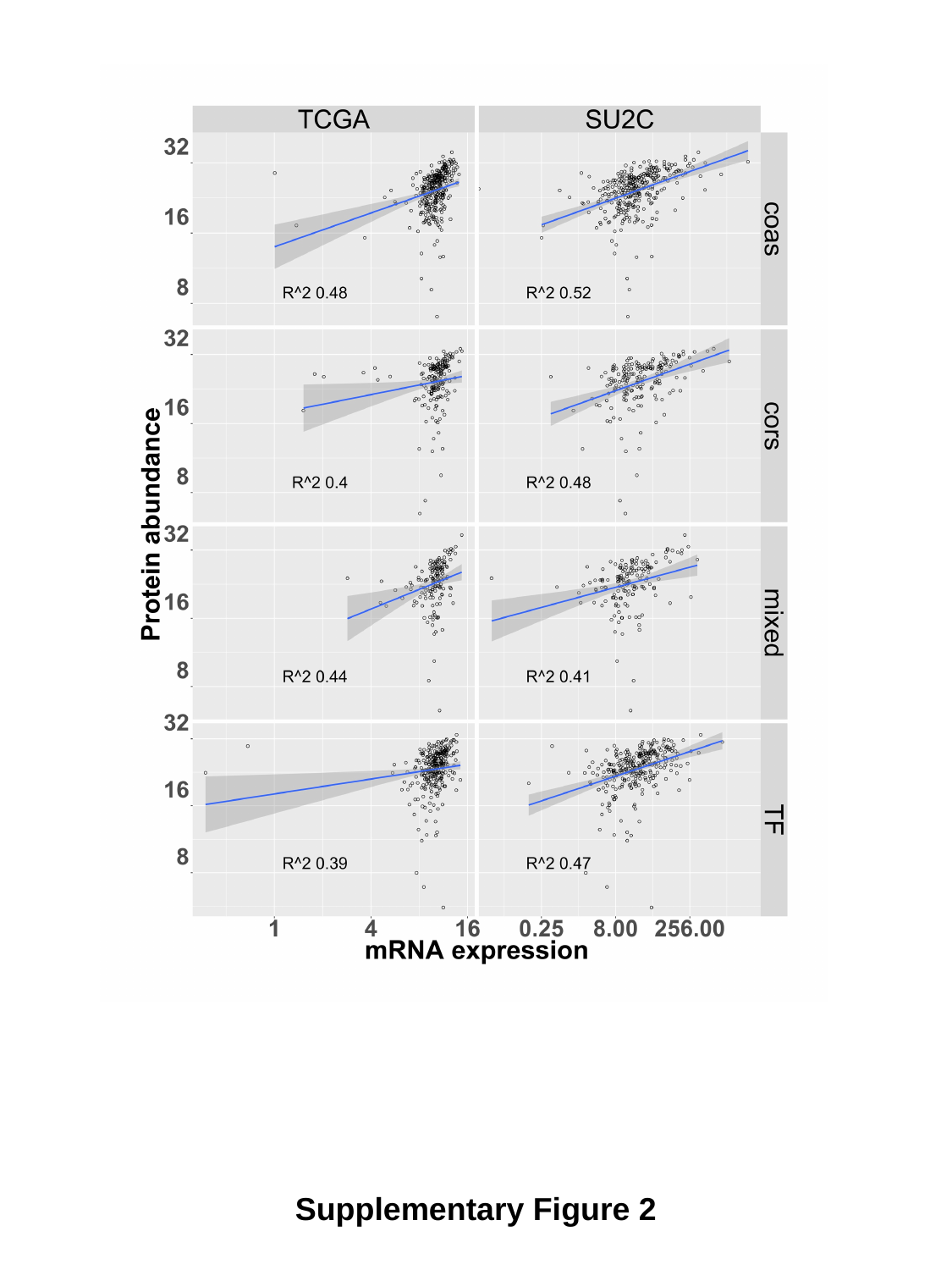

Supplementary Figure 2

### Slide 3
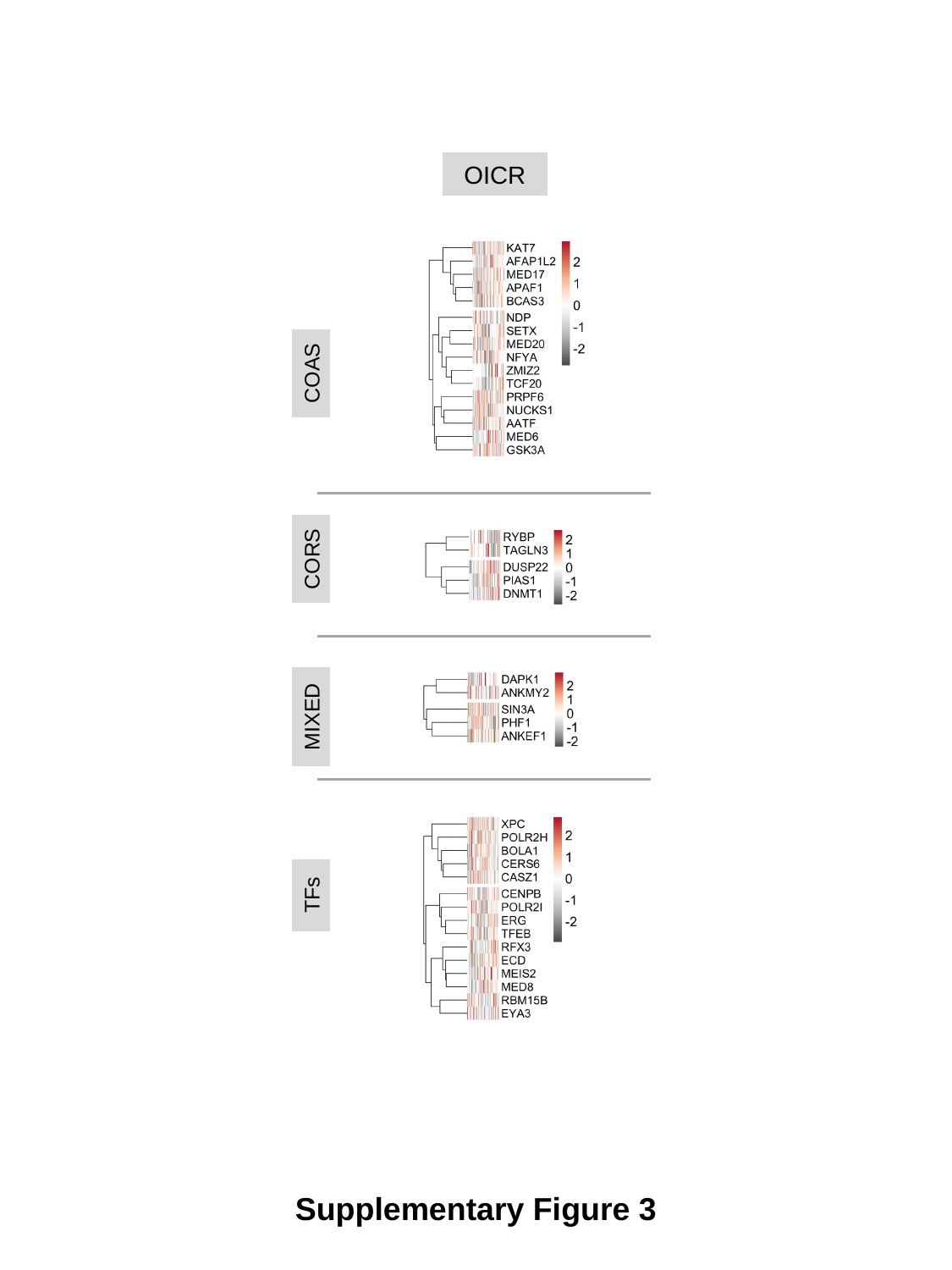

OICR
COAS
CORS
MIXED
TFs
Supplementary Figure 3

### Slide 4
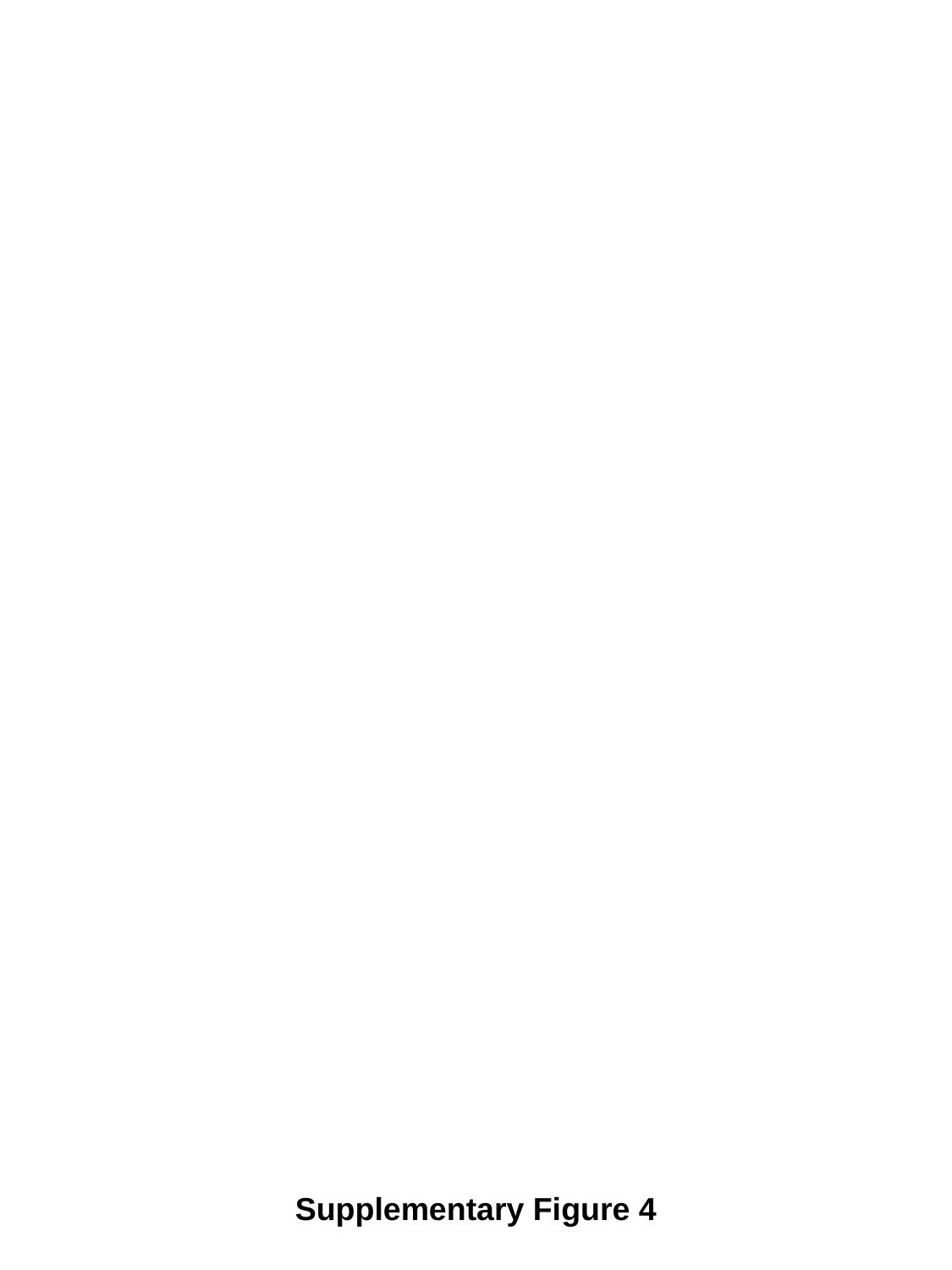

Supplementary Figure 4

### Slide 5
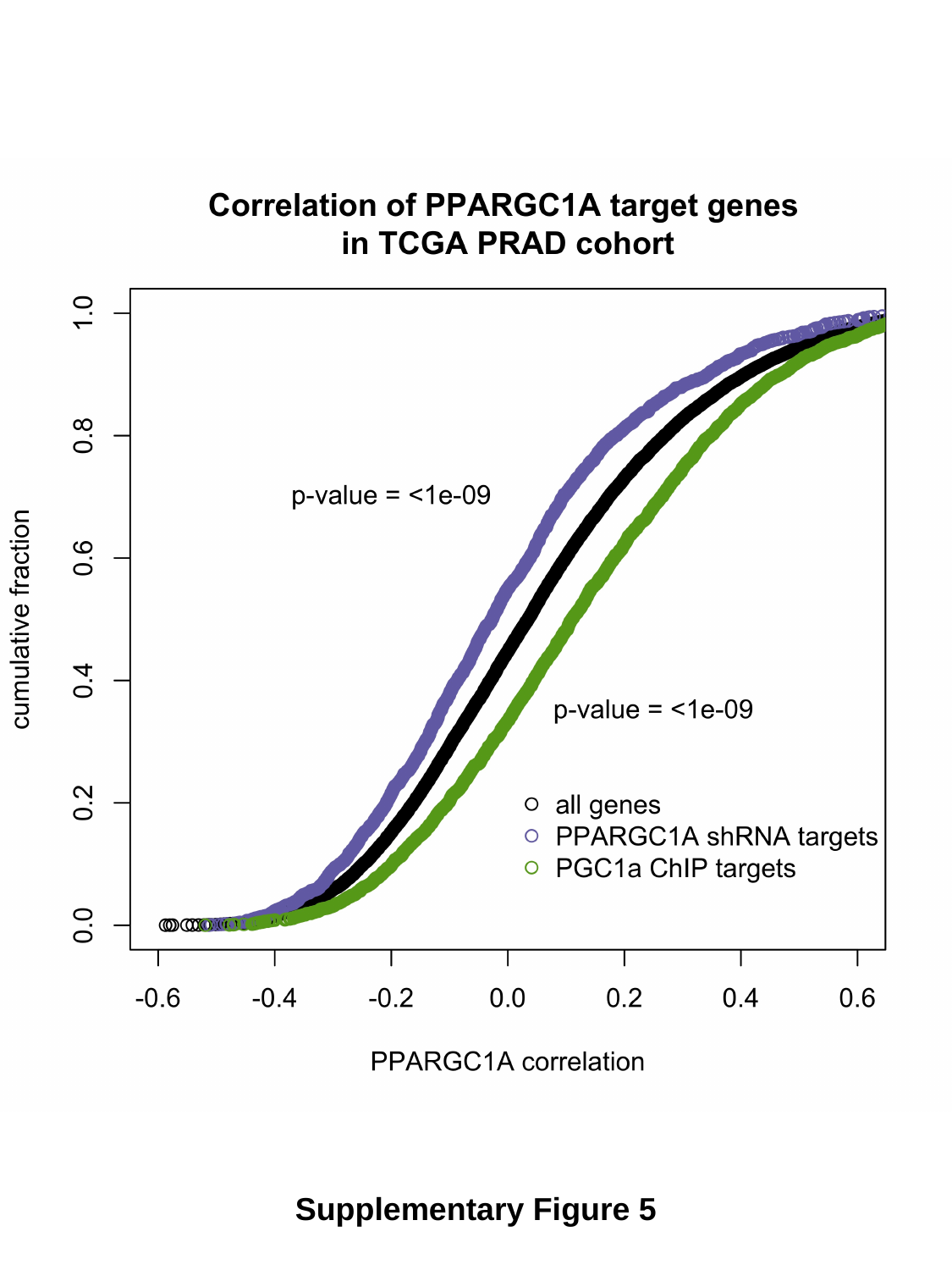

Supplementary Figure 5
